# ENVIRONMENTAL PERTURBATIONS AND TRANSITIONS BETWEEN ECOLOGICAL AND EVOLUTIONARY EQUILIBRIA: AN ECO-EVOLUTIONARY FEEDBACK FRAMEWORK

**DOI:** 10.1101/509067

**Authors:** Tim Coulson

**Affiliations:** Department of Zoology, Mansfield Road, University of Oxford, Oxford, OX1 3SZ, UK

**Keywords:** body size evolution, dynamic energy budgets, global environmental change, integral projection models, life history theory, paradox of stasis, quantitative genetics, stochastic demography, Trinidadian guppy

## Abstract

I provide a general framework for linking ecology and evolution. I start from the fact that individuals require energy, trace molecules, water, and mates to survive and reproduce, and that phenotypic resource accrual traits determine an individual’s ability to detect and acquire these resources. Optimum resource accrual traits, and their values, are determined by the dynamics of resources, aspects of the environment that hinder resource detection and acquisition by imposing risks of mortality and reproductive failure, and the energetic costs of developing and maintaining the traits – part of an individual’s energy budget. These budgets also describe how individuals utilize energy by partitioning it into maintenance, development and/or reproduction at each age and size, age and size at sexual maturity, and the size and number of offspring produced at each reproductive event. The optimum energy budget is consequently determined by the optimum life history strategy that describes how resources are utilized to maximize fitness by trading off investments in maintenance, development, and reproductive output at each age and size. The optimum life history in turn determines body size. An eco-evolutionary feedback loop occurs when resource accrual traits evolve to impact the quality and quantity of resources that individuals accrue, resulting in a new optimum life history strategy and energy budget required to deliver it, a change in body size, and altered population dynamics that, in turn, impact the resource base. These feedback loops can be complex, but can be studied by examining the eco-evolutionary journey of communities from one equilibrium state to another following a perturbation to the environment.

## Introduction

In the lowland streams of Northern Trinidad, guppies (*Poecilia reticulata*) live in complex, stable, communities (Reznick and Endler 1982). They predominantly feed on abundant stream invertebrates of high nutritional quality that are also food for a number of other fish species. Guppies themselves are prey for predatory fishes, such as wolf fish (*Hoplias malabaricus*) and pike cichlids (*Crenicichla frenata).* Predation is the primary cause of death for guppies living in these communities, such that their populations are limited by predation. In response to this limitation, guppies have evolved streamlined bauplans, live in shoals, have high maximum swimming speeds, and dull coloration to aid with the acquisition of resources in a predator-rich environment. They have also evolved fast metabolic rates, small sizes and low ages at sexual maturity, large litters of small offspring, and fast life histories to rapidly utilize the energy accrued (Reznick and Endler 1982, Reznick and Bryga 1987, Reznick and Yang 1993, Reznick et al. 2001, Travis et al. 2014). Evolution has consequently enabled these guppies to optimally detect, acquire, and utilize food in an environment where the risk of death in the jaws of a predator is near constant and high.

Above waterfalls, on the same streams where high-predation guppies are found, predatory fish are largely absent. Guppies in these streams live at high density in simpler, stable, communities, with only one major competitor, Hart’s killifish (*Rivulus hartii*), which is also an intra-guild predator. The guppy populations are food-limited, the limiting resource is therefore food availability, and the primary cause of death and failure to breed is starvation. A consequence of this is that high-quality invertebrates are rapidly consumed and are largely absent, so that guppy diets contain a high percentage of low-quality food such as algae and bacteria, that are largely absent from the diets of high-predation guppies. Guppies adapted to these low predation environments have evolved brighter male coloration, a less-streamlined bauplan, reduced shoaling behaviour, altered mouth morphology, a lower metabolic rate, greater age and larger size at sexual maturity, smaller litter sizes, larger offspring and a slower life history (Reznick and Endler 1982, Reznick and Bryga 1987, Reznick and Yang 1993, Reznick et al. 2001, Travis et al. 2014). These traits have evolved to enable guppies to detect, acquire, and utilize food in a relatively safe environment compared to the high predation one, but where the major challenge is securing enough resources to thrive.

When high predation guppies are introduced to low predation environments, their population increases in size, the age, sex and phenotypic trait structure of the population changes, and they rapidly evolve, with the community moving from the high predation equilibrium to the low predation one. Because these changes are repeated across streams, they are predictable in that we know what will happen when guppies are released from predation (Reznick and Endler 1982, Reznick and Bryga 1987, Reznick and Yang 1993, Reznick et al. 2001, Travis et al. 2014).

What characterises the Trinidadian freshwater ecosystems when they are at ecological and evolutionary (quasi)-equilibrium in either the high- or low-predation state? The size and genetic-, phenotypic trait-, size-, age-, and sex-structure of all populations in the community show no persistent temporal trends with time. I consequently use the term ‘equilibrium’ to describe a stochastic but stationary state of populations and the community. This means there may be temporal fluctuations in the sizes and structures of populations within the community, but none will exhibit a long-term temporal trend (Lundberg, Ranta, and Kaitala 2000). I use the prefix ‘quasi’ to acknowledge that some slow temporal patterns in genotype frequencies may occur in cases of slow co-evolution between species, but, that in general, natural systems frequently achieve states where statistics describing each population within the community remain within bounds for protracted periods of time (Figure 1; (Lewontin 1969, Bronstein et al. 2004). A consequence of this is there will also be no persistent temporal trends in community size and structure, or energy and nutrient flows.

**Figure 1.**
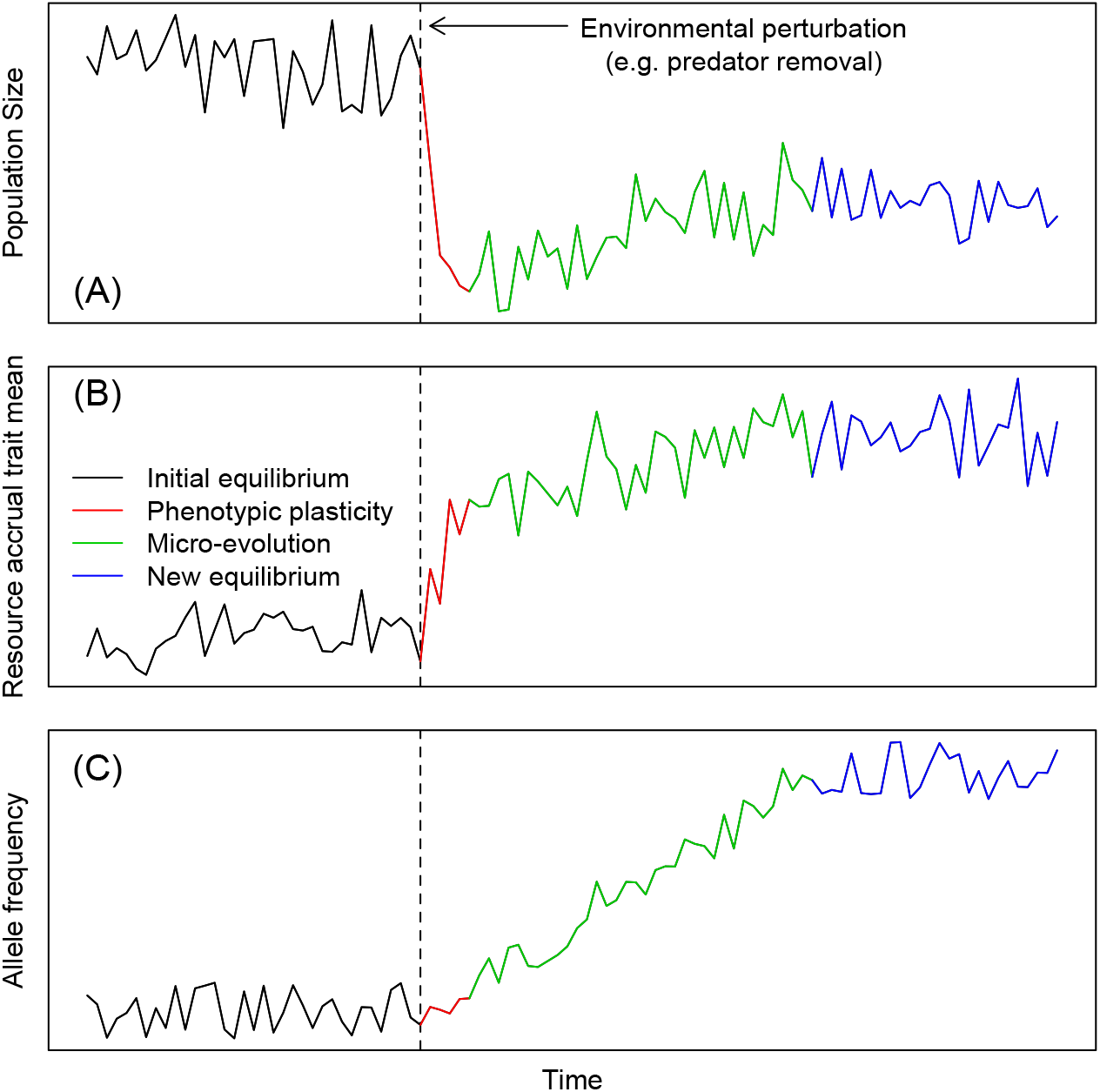
The (A) population, (B) phenotypic trait, and (C) micro-evolutionary dynamics of populations as they move from one stationary state to another. At ecological and evolutionary equilibrium, there are no temporal trends in population size, phenotypic trait distributions, or allele frequencies (black lines). Following an environmental perturbation (dotted vertical line) ecological and evolutionary change occurs (red and green lines) until a new stationary state is achieved (blue lines). In this example, population size changes quickly following the perturbation, while phenotypic and evolutionary changes occur more slowly. There will be a period when phenotypic plasticity is likely stronger than microevolution, which is reflected in quite rapid phenotypic change (B). Evolution occurs more slowly, resulting in gradual change in population size, the mean of phenotypic traits, and allele frequencies. I have given an example of ecological and evolutionary change in a single population, but similar patterns will be observed in other species within the community.

The existence of a (quasi)-equilibrium state raises a paradox. When systems are in (quasi)-equilibrium states, all heritable phenotypic traits in all species are expected to be at, or very close to, their optimal values, as otherwise phenotypic evolution would be expected to occur and the system would not be at an evolutionary equilibrium. When I refer to phenotypic traits, I define them as any attribute that can be characterised at the individual level, ranging from molecules within cells to lifetime reproductive success. In the wild, many traits including measures of body size, lifetime reproductive success, and those that describe the timing of events such as birth, are under directional selection, are heritable, but do not evolve as predicted (Merilä et al. 2001a, Estes and Arnold 2007, Haller and Hendry 2014, Rollinson and Rowe 2015, Bonamour et al. 2017). Instead they exhibit stasis that is consistent with the system being at equilibrium. The paradox is why do we predict phenotypic evolution but not see it? I address this paradox later in the paper.

An environmental perturbation, such as the loss of a predator species, leads to a period of transience within a community as population sizes and structures change (Hastings 2004, Georgelin et al. 2015). In time, the system will achieve a new (quasi)-equilibrium state (Figure 1) (Brown and Vincent 1992, Lundberg et al. 2000, Johansson and Dieckmann 2009) The observation of almost immediate evolution following a biotic or abiotic shift raises a second paradox. Evolution requires additive genetic variation within a population, but if phenotypic traits are at their optimum values, they are subject to stabilising selection, which is expected to erode additive genetic variation (Crow and Kimura 1970). The evolutionary equilibrium attained by a system should consequently exhibit low levels of additive genetic variance. Yet populations in ecosystems in the wild in (quasi)-equilibrium states express substantial amounts of additive genetic variance for many traits (Crow and Kimura 1970, Kruuk et al. 2000). The maintenance of genetic variance is a second paradox that requires a solution if we are to understand why perturbed systems move from one evolutionary equilibrium to another. I also consider this paradox later in the paper.

The genetic diversity paradox has similarities with another problem that has long interested biologists – why do so many species coexist when they appear to share niches and rely upon the same resources (Chesson 1986, Wright 2002)? The coexistence problem is also relevant to the problems I address in this paper, and in particular why the ecological and evolutionary (quasi)-equilibria contain so many coexisting species. I return to this topic later in the paper.

Before reviewing properties of ecological and evolutionary equilibria, and considering ways to predict the ecological and evolutionary transients between contrasting stationary states following a biotic or abiotic change, I do two things. First, I consider how the different levels of biological organisation – alleles, genotypes, phenotypes – are linked by a number of fundamental processes (Figure 2) (Coulson et al. 2017).

**Figure 2.**
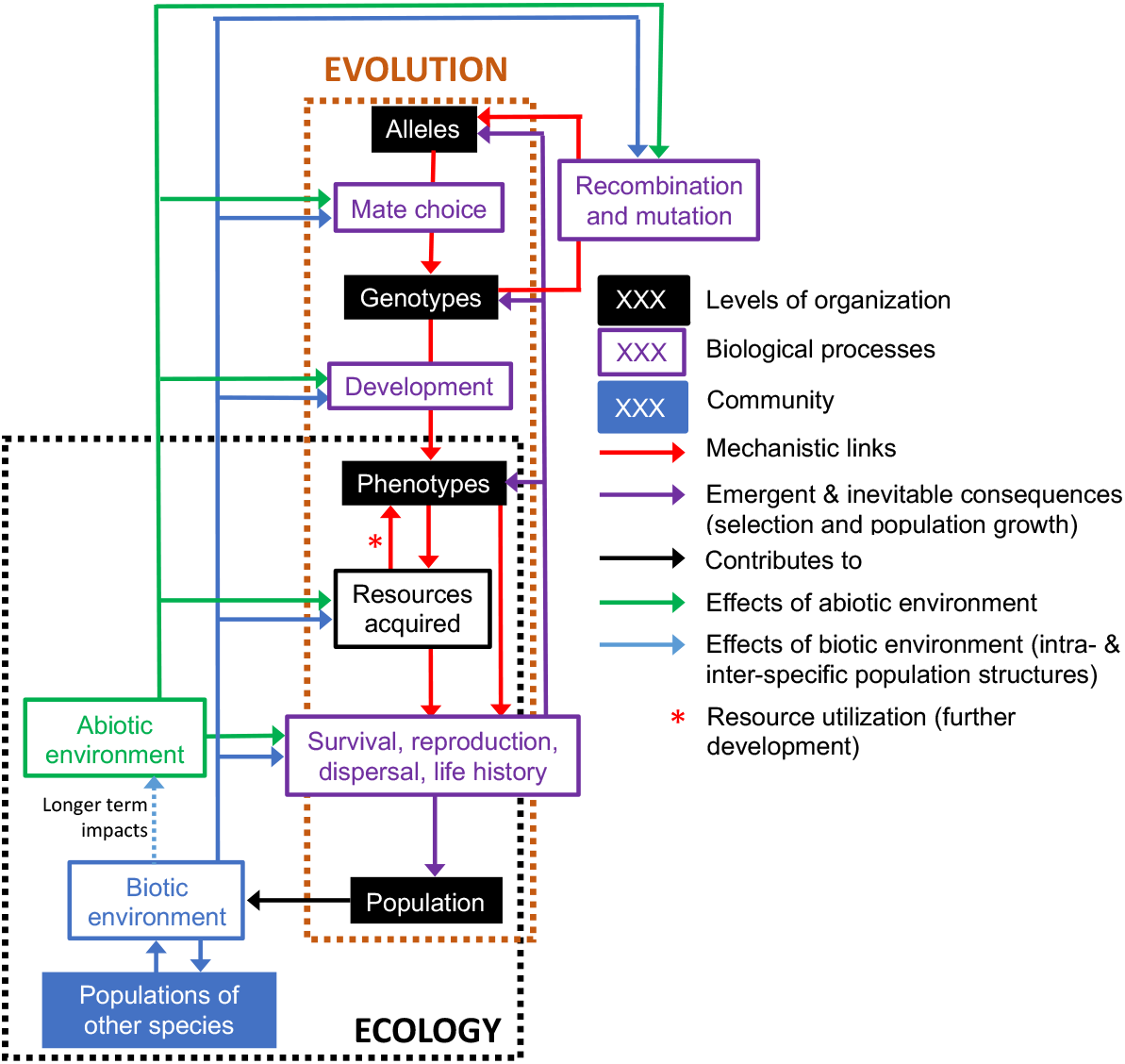
Complex eco-evolutionary feedbacks can arise from the dynamics of distributions of alleles, genotypes and phenotypes (levels of biological organization). These levels are mechanistically linked by key biological processes. Mate choice determines how alleles form genotypes in offspring, with development determining how phenotypes develop from genotypes. Phenotypes impact which resources individuals acquire and can influence survival, reproduction, dispersal and life history (demography). Acquired resources, such as a territory, can also influence demography directly, while also feeding back to impact further phenotypic development (including state-dependent behaviors). Selection and population dynamics are emergent consequences of which phenotypes survive and reproduce. The size and structure of the focal species and those species with which it interacts, constitute the biotic environment. The biotic environment, along with the abiotic environment, can directly influence the distribution of resources, but also biological processes in both the focal species and other species within the community. Recombination and mutation generate genetic variation and can be influenced by the environment.

Parental alleles combine, through patterns of mate choice, to generate the distribution of offspring genotypes, which in turn develop into a distribution of offspring phenotypic traits. These traits determine individual abilities to acquire resources from the ecological environment in which they find themselves, which in turn determines continued rates of development and chances of survival, reproduction and dispersal at each age and size. Both the population dynamics, and the strength of selection that can alter the distribution of phenotypic traits, genotypes and alleles, are inevitable emergent properties of these rates. Recombination and mutation determine the distribution of alleles among individuals selected to be parents (Crow and Kimura 1970). The fundamental biological processes of mate choice; development; survival, reproduction, and dispersal; and mutation, can all be influenced by aspects of the biotic and abiotic environment. The biotic environment experienced by an individual consists of the numbers and attributes of all individuals it interacts with whether they are con- or hetero-specifics. These interacting individuals include its parents, who can use their attributes to influence offspring phenotypes via controlling the developmental environment.

Second, I introduce the concept of resource accrual traits. These are the set of traits associated with the detection and acquisition of resources required for life (Abrams and Chen 2002): energy, trace molecules and water required for survival, and mates necessary for reproduction (at least in sexually reproducing diploid species) (Brown et al. 2004). Within the set of resource accrual traits are resource detection traits such as the size and structure of sense organs and the brain, and resource acquisition traits such as foraging behaviours and aspects of morphology. I will argue that these traits are likely to be subject to stabilising selection, yet that individual heterogeneity in the types of resources individuals specialise on (Bolnick et al. 2007), act to maintain additive genetic variation underpinning these traits. Desirable resource accrual traits, and their optimum values, are specific to the environment in which individuals find themselves. For example, the traits expressed by high-predation guppies in high-predation environments (Reznick and Endler 1982) are well-adapted to an environment where the limiting factor is predation and the primary cause of death is predation. In contrast, in the low-predation environments, guppies express traits that make them well-adapted to an environment where the primary cause of death is starvation and the population is food-limited.

In addition to optimising resource accrual traits, evolution also optimizes energy utilisation to maximise fitness (Kooijman and Kooijman 2010). It does this by optimising the life history via evolving an energy budget to optimally allocate energy to maintenance, development, and reproduction, at each age and size (Hin and de Roos 2019). The energy budget determines age and size at sexual maturity, the number and size of offspring produced at each reproductive attempt, and life expectancy (Stearns 1976).

I primarily focus on the guppies as they provide a well-characterised example of a general phenomenon: if the primary causes of death or failures to breed in a population are altered by a biotic or abiotic perturbation, so too is the limiting factor, the population dynamics (Schindler 1974, Sibly and Hone 2002, Rohr et al. 2003), selection, and the course of evolution (Walters and Juanes 1993, Fishman and Willis 2008). Such a perturbation results in rapid change in population size and selective regimes in a focal species and the species with which it interacts. The new regime selects for phenotypic traits and trait values that allow organisms to optimally detect and acquire energy, and a life history that allows these resources to be optimally utilised given the primary causes of death and failures to breed in the new environment in which they find themselves. These processes determine the transient dynamics between ecological and evolutionary (quasi)-equilibria.

In the remainder of this paper I provide 1) a description of the conditions required for a system to be at ecological and evolutionary equilibria, 2) discussion of how these conditions can be achieved in nature, and the observation of two paradoxes that need to be addressed: the paradox of stasis for size-related phenotypic traits, and the paradox of standing additive genetic variation, 3) solutions to the paradoxes, 4) a more detailed treatment of the evolution of life history and body size, which I consider to be central to solving the paradox of stasis, 5) an examination of models that can describe systems in ecological and evolutionary equilibrium, 6) a focus on how systems transition between different equilibria states as a consequence of abrupt environmental change, and 7) a brief concluding section.

## Characterising populations at ecological and evolutionary equilibria

I start by considering a single population. I define a population as being stationary and at a stochastic ecological and evolutionary equilibrium when both its size and genetic, phenotypic, sex, and age structure fluctuate within bounds but show no persistent temporal trends (Caswell et al. 2004). In formal demography, this equilibrium is a special case, as populations can achieve structural stationarity, yet can also change in size with time as the population grows at a constant stochastic rate (Tuljapurkar 1989). Because I require a stationary distribution of population sizes *N_t_*, the special case I consider requires the long-run stochastic growth rate *a* to equal zero, where

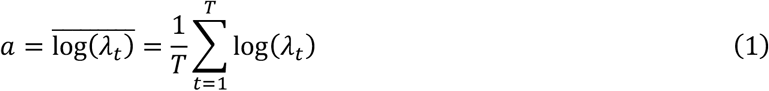

with *t* being time (Tuljapurkar and Orzack 1980). The equation defines the long-run stochastic growth rate of the population as the arithmetic mean of the logged per-time step population growth rates *λ_t_* calculated across a time series of length *T* where 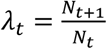. But how does this formulation relate to the ecological and evolutionary equilibria I describe above, where there is also no persistent temporal trend in allele and genotype frequencies, and phenotypic trait means and variances? Examining the determinants of *λ_t_* helps answer this question.

I consider a population as consisting of individuals, each with a very large number of phenotypic trait and genotypic attributes. From these individuals, I can construct a multivariate distribution describing the number of individuals in each individual attribute (or structural) class at time *t* (Ellner et al. 2016). For example, in a very simple case where a population is structured by the individual attributes sex, age, weight, and genotype at a locus *X*, there may be 23 individuals within the population at a particular point in time that are male, aged 35 years, weigh 82kg and express genotype *AA* at locus *X*, but only seven individuals that are female, aged 74, weigh 52kg with genotype *BB* at locus *X*. All other sex, age, weight and genotype classes will also contain a particular number of individuals (that could be zero). Over time, the numbers of individuals in each structural class can fluctuate (Ellner and Rees 2006), but as long as there are no persistent temporal trends in these time series of numbers, the population will be at a stationary equilibrium. It is possible to estimate the long-run stochastic growth rate for each structural class using equation (1) above; at ecological and evolutionary equilibrium, these rates will all be zero (Tuljapurkar and Orzack 1980).

The per-time step growth rate of the entire population 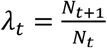 can be expressed as *λ_t_* = *S_t_* + *R_t_* where *S_t_* is the annual per capita survival rate, and *R_t_* is the annual per capita recruitment rate (defined as the per capita production of offspring that are produced between times *t* and *t* + 1 that survive to enter the population at time *t* + 1 weighted by the relatedness of offspring to parents (Coulson et al. 2006)). If we ignore dispersal, change in population size over a time step is consequently caused by any difference in the numbers of individuals that enter the population via birth, and the numbers that leave it via death. But what determines *S_t_* and *R_t_* at a particular time *t*? Both rates vary across structural classes (e.g. individual attributes), and with the environment (Leslie 1945, Caswell 2001). The values of *S_t_* and *R_t_* at any particular time t consequently depend upon the population’s genetic, phenotypic trait, age and sex structure.

Survival, development, reproduction, mutation, recombination, mate choice, and genetic and non-genetic inheritance are all processes that can alter the population’s structure between time *t* to *t* + 1 (Coulson et al. 2017). With the exception of genetic inheritance, all these processes can be influenced by aspects of the biotic or abiotic environment, and it is environmental variation that continually generates variation in population structure via impacting these processes (Figure 2). Structured demographic models based on Figure 2 can be constructed to capture the effects of the environment on each of these processes, and used to iterate distributions of population structure and size from one-time step to the next (Coulson et al. 2017).

Structured demographic models have been constructed for small numbers of individual attributes such as one or two genetic loci (Charlesworth 1994), one or two phenotypic traits (Easterling et al. 2000), age (Leslie 1945), or sex (Schindler et al. 2015). Despite a focus on only a small number of attributes, these models have provided significant ecological and evolutionary insight. Models consist of functions that capture the processes (Figure 2) that can alter distributions of individual attributes by moving individuals into and out of each structural class at a point in time (Ellner et al. 2016). For example, survival functions determine the proportion of individuals within a class that remain within a population, development functions determine the rates at which surviving individuals move between structural classes, reproductive functions determine the expected number of offspring produced by individuals within each structural class, while mutation, mate choice, and genetic and non-genetic inheritance functions, determine the attributes of offspring and the structural classes in which they enter the population. I cumulatively refer to survival and reproductive functions as demographic functions, and mutation, mate choice, and genetic and non-genetic inheritance functions as inheritance functions. Any function, with the exception of those that describe genetic inheritance, can contain terms describing the influence of biotic and abiotic environmental drivers (see section **Population models of ecological and evolutionary stasis** for additional details).

Structured demographic models do not need to include all levels of biological organization found within a population (Figure 2). For example, models have been constructed that structure populations by only age, genotype, or phenotypic trait, or by just two of these three levels of organization. When these simple models are constructed, they make assumptions about the processes that link levels of biological organization that are not explicitly incorporated into models (Traill et al. 2014). For example, models that include genotypes, but not phenotypes, assume that each genotype always develops with perfect fidelity into the same phenotype trait value, such that fitness can be directly predicted from the genotype without having to model phenotypic development (Charlesworth 1994). In this paper I primarily focus on cases where this assumption is not made, and populations are structured by alleles, genotypes and phenotypic traits (Coulson et al. 2011, Coulson et al. 2017).

As previously mentioned, each genotype (and consequently allele) will have a long run stochastic growth rate of zero at ecological and evolutionary equilibrium. There is another equivalence that individuals in particular structural classes will share: when genotypes are additive, and in the absence of mutation, genotypes within offspring cohorts are expected to have identical reproductive values (Sæther and Engen 2015). The expected reproductive value is the expected number of descendants of an individual in a particular structural class at time t, at some distant time point in the future (Fisher 1930, Caswell 2001). There are two ways expected reproductive value can be calculated for genotypes from a model. First, by keeping track of the number of descendent copies of each allele within the population with time (Barton and Etheridge 2011). Second, by recording the number and relatedness of descendants alive in future time steps of individuals that carried a particular allele at time t (Fisher 1930). Descendants are included in the calculation regardless of whether they carry alleles found in the original genotype themselves; genetic inheritance in sexually reproducing species means that descendants do not necessarily carry ancestral alleles. The two quantities are expected to be equivalent, although stochasticity in survival, reproduction and inheritance can result in an individual with a particular allele having very many descendants with none, or very few, carrying copies of it. Reproductive value can be calculated for any structural class – genotype, phenotype, age, sex. However, at evolutionary equilibria, reproductive values of offspring genotype or phenotypic trait classes are only equivalent when genotypes and phenotypes are additive functions of underlying alleles. This requires a little more explanation.

In structured models of sexually reproducing species, alleles do not form structural classes. Instead, these classes consist of genotypes or phenotypic traits (Charlesworth 1994, Childs et al. 2016, Coulson et al. 2017). The dynamics of alleles, and hence microevolution, can be determined from the dynamics of the number of individuals within genotypic classes. A genotype is considered additive, if its contribution to a phenotypic trait is determined solely by the sum of the contribution of each of its alleles to a phenotypic trait (Falconer 1960). If a phenotypic trait value is determined by an interaction between alleles at a locus (dominance) or by interactions between genotypes at different loci (epistasis), the assumption of additivity is violated. A consequence of this is that microevolution cannot be inferred from the dynamics of phenotypic trait values.

If all genotypes at all loci that contribute to a phenotypic trait are additive, then the dynamics of the phenotypic trait, the genotypes that contribute to the trait, and the alleles that constitute the genotypes, are asymptotically predicted by their reproductive values (Fisher 1930). Under these circumstances, when different genotypes and phenotypic trait values expressed in offspring cohorts have different reproductive values, then microevolution will occur and allele frequencies will change. When they all have identical reproductive values, the population is at evolutionary equilibrium.

Expected reproductive value is a key concept in biology as it is the expected asymptotic fitness of an individual with particular attributes (Fisher 1930). The reproductive value of a structural class is determined by the life history of individuals with those attributes. A life history describes key events within life – age and size at sexual maturity, the timing of reproductive events, the size and number of offspring produced at each successful reproductive attempt, and the age and size at death (Stearns 1976). Life history strategies can consequently be characterized with size- and age-specific survivorship, development, fertility and inheritance schedules (Figure 3) (Stearns 1976) – all of which can be calculated from the functions that constitute structured demographic models (Childs et al. 2004).

**Figure 3.**
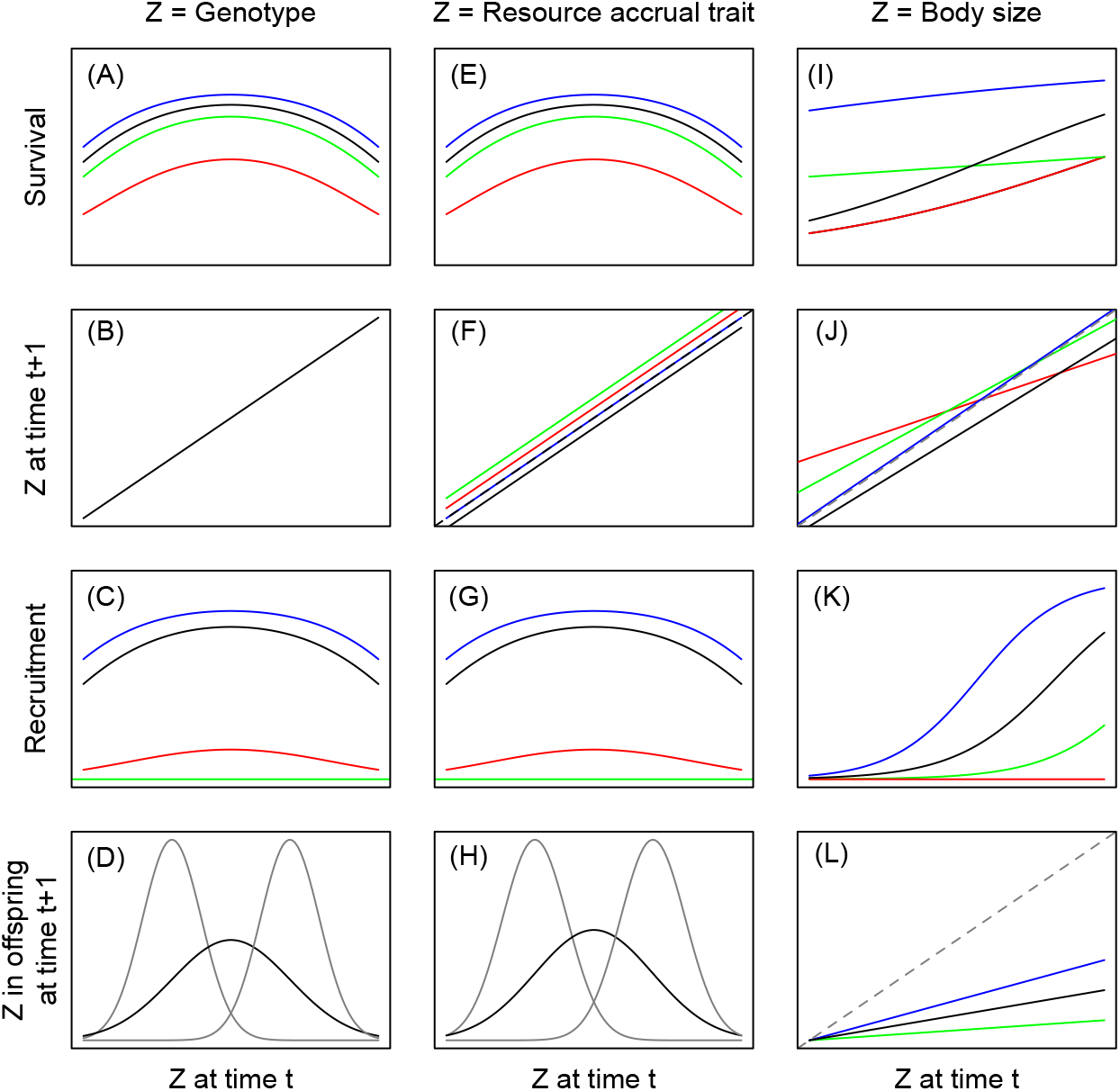
The form of survival (A, E, I), development (B, F, J), reproduction (C, G, K) and inheritance (D, H, L) functions for different individual attribute types. (A-D) form of functions for a breeding value or genotype that is subject to stabilizing viability (A) and fertility (C) selection, that does not develop throughout life (B), and exhibits genetic inheritance (D). The two narrower grey distributions in (D) represent breeding values or genotypes in parents, while the flatter black distribution represents the distribution in offspring following genetic inheritance. Similar dynamics are observed for a resource accrual trait that maintains its shape throughout life but increases in size with age as the environmental component of the phenotype develops (F). The trait is genetically inherited (H). Body size (I-L) is subject to directional viability (I) and fertility (K) selection, develops with age, and appears heritable (L) with parents and offspring have more similar values than non-related individuals. Different colored lines represent different ages, showing how functions can vary with age. In (B), (D) and (H) the same functions are observed at each age.

A life history strategy plays out on a per generation time scale, and can be described with per generation summary statistics such as generation length, mean lifetime reproductive success *R*_0_, and life expectancy (Caswell 2001, Tuljapurkar et al. 2009a, Steiner et al. 2014). *R*_0_ describes the rate of population growth per generation and is a per generation version of *λ_t_*. In a deterministic population that exhibits no temporal trend in numbers log(*R*_0_) = 0.

*R*_0_ can be calculated for individuals within each genotypic or phenotypic structural class (Tuljapurkar et al. 2009b). When additivity is assumed, individuals in all structural classes will have equal expected values of *R*_0_ when the population is at ecological and evolutionary equilibria. When the assumption of additivity is violated, this will not necessarily be the case.

Biologists are often interested in the evolution of life histories, without focusing on evolution of the resource accrual traits that influence energy acquisition and the optimal life history (Stearns 1976). To do this, life history strategies are assumed to be clonally and near perfectly inherited, rather than as arising as a consequence of underlying resource accrual traits (Childs et al. 2004). Under this assumption, the reproductive value of a life history will be associated with its long-run stochastic growth rate when measured in the presence of competing, clonally inherited, life histories. *R*_0_ can usually (but not always) also predict which clonally inherited life history will grow to dominate all others. Such approaches rely on the assumption of clonal inheritance. The approach is useful as invasion approaches can be used to identify the evolutionarily stable life history strategy, and in particular the optimal value of life history trade-offs such as those between offspring size and offspring number. However, genetic architecture significantly influences ecological and evolutionary dynamics (Schreiber et al. 2018), except in very specific cases (Geritz and Éva 2000, Metz and de Kovel 2013), and I consequently focus on cases where inheritance is not assumed to be clonal.

The conditions for a stochastic ecological and evolutionary equilibrium within a population – a long-run stochastic growth rate of zero for each structural class – are relatively straightforward, but they impose constraints on the values of reproductive value and the life history when additivity is assumed. Structured demographic models of populations at stochastic ecological and evolutionary equilibrium can be constructed from demographic, development, and inheritance functions that describe the rates of movement of individuals between structural classes from time t and t + 1. From these models, it is then possible to calculate long-run stochastic growth rates, reproductive values, temporal dynamics of the number of individuals, and descriptors of the life history for each class (Coulson et al. 2011, Coulson et al. 2017).

Structured demographic models provide a remarkably powerful tool to study the dynamics of populations, allele and genotype frequencies, and phenotypic trait distributions. The properties of models at ecological and evolutionary equilibria are also well understood. However, for these equilibria to be achieved, particular conditions are necessary.

## Conditions required for ecological and evolutionary stationarity

A long-run stochastic growth rate of *a* = 0 in a single species population requires density-dependence: a pattern that describes a decline in the per time step population growth rate *λ_t_* as population size increases (Engen and Sæther 1998). Density-dependence occurs when a direct species interaction, such as predation, herbivory, or rarity of a resource species, limits a population’s growth rate (Hassell 1975). Populations are consequently limited by aspects of the environment that prevent organisms detecting, acquiring and utilizing sufficient resources to survive and reproduce (Murray 1982). Limiting factors determine key causes of mortality or failures to breed within a population (Stearns 1976, Gaillard et al. 2000). For example, prey species that are predator-limited are limited by their ability to accrue resources while avoiding predation.

Ecologists have long been interested in how species that rely upon the same resources or limiting factor can coexist (Gause 1932). The general solution to this problem is that coexistence of competing species occurs when individuals within a species compete more strongly for a resource than individuals between species. In contrast, if interspecific competition is stronger than intraspecific competition, then the better competitor species will drive the other extinct (MacArthur and Levins 1967).

Four different mechanisms can contribute to the relative strengths of intra-(density dependence) and interspecific competition for a resource or limiting factor. They act by either increasing the strength of intraspecific competition or by reducing the strength of interspecific competition (Chesson 2000b). The first mechanism, termed density-fitness covariance, does the former. It describes any mechanism that acts to increase the strength of intraspecific competition relative to interspecific competition, such that each population increases in density when its numbers drop (Chesson 2000a). An example of this is when competing species utilise the same resource, but are segregated in space, such that they tend to compete with conspecifics rather than with hetero-specifics. The second set of mechanisms is termed “stabilising” and works by decreasing the strength of interspecific competition by ensuring each species has a refuge that protects it from extinction (MacArthur and Levins 1967). An example of a stabilising mechanism is two species that do not have completely overlapping niches, but instead partition some resources or limiting factors between themselves. The third coexistence mechanism is termed the storage effect, which occurs when different species respond to environmental variation in space or time in contrasting ways, with different species being more competitive in some types of environment than others (Warner and Chesson 1985). For example, one species may be a better competitor in winter, while the other may gain an advantage in summer. The final mechanism is termed relative non-linearity where different species benefit in different ways from fluctuations in the abundance of the limiting resource via non-linear effects of environmental variation on their population growth rates (Chesson 1994). These last two mechanisms can contribute to coexistence when they act to increase the strength of intraspecific competition to a greater extent than they increase the strength of interspecific competition.

To incorporate these ecological coexistence mechanisms into demographic models it is necessary to construct a model for each interacting species, and to couple them (Adler et al. 2010, Bassar et al. 2017). This can be achieved by making the demographic, development, or inheritance functions of each species a function of the size or structure of both the focal species and of the competitor species. Such models capture the indirect effects of two species competing for a resource or other limiting factor; the resource or shared predator are not explicitly included in models. Direct effects can be incorporated into models by formulating demographic, development, and inheritance functions of focal species to be functions of the size and structure of interacting resource or consumer species on neighbouring trophic levels (Lachish et al. in press).

Ecological coexistence of multiple species requires direct interactions to limit each population such that its long-run stochastic growth rate is zero. This might sound like a big ask, but most species only interact strongly with a relatively small number of others, and when this happens, interacting species often coexist by limiting one another (Allesina and Tang 2012). Consider, for example, the case of the lynx and snowshoe hare, where both species cycle, with the predator dynamics lagging a quarter of a cycle behind that of the prey (Keith and Windberg 1978), a pattern often termed Lotka-Volterra predator-prey dynamics. The predatory lynx is limited by availability of the snowshoe hare prey, particularly in periods following high lynx abundances. In contrast, the hare is limited by lynx predation, particularly when lynx are abundant. The strength of these limiting factors fluctuate (reasonably) predictably with time, and the two populations cycle. Their long-run stochastic growth rates are both zero, but the numbers of both species cycle, in an endless entwined pattern, where each species limits the other, at least some of the time (Krebs et al. 1986). Because many limiting factors are biotic, it appears that ecosystem stability may be the norm, particularly when limitation of each population operates primarily via a small number of strongly interacting species.

The case of the lynx and snowshoe hare is a special one, and few systems are this simple. However, different species that share a resource or limiting factor are able to coexist via one or more of the mechanisms described above because they have adaptations that allow them to do so. In particular, different species may have adaptations that increase their fitness by making them specialists in particular environments (e.g. the storage effect) or by allowing them to utilise a unique resource that is unavailable to individuals of competitor species (e.g. a stabilising mechanism) (MacArthur and Levins 1967, Warner and Chesson 1985). Successful species are often those that have been able to innovate in a way that allows them to utilise a previously unobtainable or under-utilised resource.

Rotifers feeding on algae provide a particularly interesting example of how phenotypic variation that allows adaptation to specific environments permits coexistence. Algae-rotifer dynamics deviate from classic Lotka-Volterra predator-prey expectations, with the predatory rotifer population lagging a half cycle, rather than the expected quarter cycle, behind that of their prey (Yoshida et al. 2003). The reason for this is that the algae come in two different phenotypic forms – clumpers and soloists (Becks et al. 2010). The two forms are clonal, and rarely reproduce sexually. They consequently act like two different species, even though they are in fact two genetically distinct clones of the same species. Individuals of the soloist clones have high fitness when the predators are rare. They can grow quickly, efficiently utilising abundant resources, and they outcompete the clumper clone. Survival is high, as rotifers are rare, and the population of soloists increases rapidly. The fitness of the soloist individuals starts to decline as predator numbers increase, and their population becomes predator limited. Rotifer numbers increase because the soloists make easy prey. However, the decline of soloists is simultaneous with an increase in the fortunes of the clumpers. Individuals of this clone band together into groups that are too large to be easily predated. They are not as efficient at turning resources into new clones as the soloists are, but when soloists are largely absence, such inefficiency is not an issue. As the clumpers dominate, the predatory rotifer population starts to decline as accessible prey abundance declines, and the time of the soloist dominance comes around once again. The two clones are able to coexist because the clumper has an adaptation that favours them in the high predation environment, while the soloists dominate in a low predation environment (Hiltunen et al. 2014). Population limitation of the two clones via resource limitation and predation allow coexistence of different phenotypes: the two clones coexist via the storage effect which is facilitated by the adaptive ability to clump or not. Mechanisms such as this must be abundant in nature give so many species coexist.

The algae-rotifer system provides a link between coexistence of two species that share a predator but cannot reproduce, and the maintenance of genetic variation in sexually reproducing diploid species. The observation is particularly useful, as it reveals that the mechanisms that allow ecological coexistence of species in communities at equilibrium, may also contribute to evolutionary equilibria – another problem that needs solving if we are to understand stationary ecosystems.

Trivial evolutionary equilibria occur when all genetic variation is eroded such that all individuals are genetically identical (Wright 1969, Crow and Kimura 1970). However, such equilibria are not informative when considering the types of evolutionary transients observed, for example, in Trinidadian freshwater streams when predators are removed and the community moves between stationary states quite rapidly (see introduction). More interesting equilibria maintain additive genetic variance. For this to happen, either stabilising selection and mutation-selection balance, a particular form of phenotypic plasticity, or frequency-dependent selection, must be in operation (Gillespie 1994). But how are these related to the coexistence mechanisms that ecologists focus on when considering species coexistence?

The strength of selection is determined by the association between phenotypic trait values and fitness (e.g. survival and reproduction) (Fisher 1930, Price 1970). It is a within generation process, most easily measured over a time step using viability and fertility differentials (Lynch and Walsh 1998). How selection translates to the underlying genotypes and alleles is determined by the genotype-phenotype map that determines how phenotypic traits develop. Directional selection requires increasing or decreasing functions between phenotypic trait values and fitness (e.g. survival and reproduction), while stabilising selection requires humped-shaped associations, where the highest survival or reproductive rates are observed at intermediate phenotypic trait values (Barton and Partridge 2000). Many evolutionary models assume simple additive genotype-phenotype maps described in the previous section that allow microevolution to be inferred directly from the dynamics of the phenotypic trait. This assumption is known as the phenotypic gambit (Lynch and Walsh 1998).

If the phenotypic gambit is assumed (and frequency-dependence is not operating), evolutionary equilibria require stabilising selection. Yet stabilising selection (as well as directional selection) erodes additive genetic variance unless some other process replaces it. When selection is weak (i.e. the hump of the association between the phenotype and fitness is quite flat), mutations can generate novel additive genetic variance at a rate that counters losses due to selection in a process termed selection-mutation balance (Bürger and Lande 1994).

Phenotypic plasticity occurs when the same genotype can generate different phenotypes in different environments (Pigliucci et al. 2006). If multiple genotypes can generate the optimal phenotypic trait value, each genotype will have maximal fitness and will be able to coexist.

The third process that can maintain additive genetic variance is frequency-dependent selection (Ayala and Campbell 1974). This occurs when a phenotypic trait value, and the underlying genotypes and alleles, experience higher survival and reproductive rates the rarer they become, such that they increase in frequency when rare. There are a number of processes that can generate frequency-dependent selection including heterozygote advantage, disassortative mate choice, spatial variation in the optimum phenotypic trait value, and heritable individual specialisation in resource use or limiting factor (Hartl et al. 1997). This last process occurs when the strength of competition increases with phenotypic similarity between competing individuals, with coexistence of individual variation being facilitated by the four coexistence mechanisms that operate at the species level (Bassar et al. 2016). For example, fitness-density covariation can occur when individuals with different phenotypic trait values feed on the same resources but in different locations – i.e. there is small-scale genetic variation underpinning phenotypic variation. Stabilising mechanisms occur when populations consist of individual specialists that exhibit different foraging phenotypes and feed on different resources (Chesson 1994). In the Cocos Island finch, some individuals exclusively forage on seeds and invertebrates found on the ground, while others forage solely on arboreal insects, rarely venturing onto the ground (Werner and Sherry 1987). When survival, reproductive, or development rates of individuals with different phenotypic trait values are impacted by environmental variation in different ways, then the storage effect can occur, and depending upon the way that this environmental variation impacts demographic, development or inheritance functions, relative non-linearity can also arise (Warner and Chesson 1985, Chesson 1994, 2000a).

The same processes can consequently maintain genetic diversity within a population, and species diversity within a community. When these processes are operating, ecological and evolutionary equilibria can be achieved, whereby each species, and each structural class in each population, has a long-run stochastic growth rate of zero. At these equilibria, a number of other properties of stationarity may be observed, including equivalence of reproductive values of genotypes in the structural classes of offspring, and equivalent mean lifetime reproductive successes of individuals regardless of their phenotypic trait values. But such patterns are not always observed, and this conveniently leads me to the paradox of stasis.

## A solution to the paradox of stasis

When the phenotypic gambit is assumed, phenotypic variation between individuals can be partitioned into a contribution attributable to additive genetic variation (Falconer 1960, Lynch and Walsh 1998). This can be estimated by statistically modelling phenotypic variation as a function of relatedness between individuals (Kruuk 2004). The ratio of the additive genetic variance of a phenotypic trait, to the variance of the trait itself, is the narrow sense heritability, *h*^2^. The closer *h*^2^ is to one, the larger the proportion of phenotypic variance that is attributable to additive genetic differences between individuals. When *h*^2^ = 0, then none of the phenotypic variation is attributable to additive genetic differences between individuals. The phenotypic variance that is not attributable to the additive genetic variance can be partitioned into various other contributions, such as dominance and epistatic variances, but the one I focus on is the environmental variance. I consequently assume that an individual’s phenotypic trait *Z_i_* is equal to the sum of an additive genetic contribution *A_i_* (termed its breeding value) and an environmental component *E_i_* such that *Z_i_* = *A_i_* + *E_i_* (Lynch and Walsh 1998).

In populations at evolutionary stasis, distributions of phenotypic traits are stationary and show no persistent temporal trends in their means or variances (Lande 2009). If additivity is assumed, and the traits are heritable (e.g. *h*^2^ > 0), this means they should not be subject to persistent directional selection (Merilä et al. 2001b, Estes and Arnold 2007, Haller and Hendry 2014, Rollinson and Rowe 2015, Bonamour et al. 2017). Paradoxically, many size-related traits, such as body mass in fish (O’Sullivan et al. 2019), and horn length and antler size in ungulates (Kruuk et al. 2002, Traill et al. 2014, Coulson et al. 2018), are heritable, under directional selection, but do not evolve as predicted (Merilä et al. 2001b). Groups of individuals with larger trait values have higher reproductive values, and greater values of *R*_0_, but they have long-run stochastic growth rates of *a* = 0. Structured demographic models that do not explicitly incorporate evolution, capture these dynamics well (Traill et al. 2014). These models predict no change in mean body size with time, even though the trait has an estimated non-zero heritability from both observation and from model predictions (Coulson et al. 2010). Put another way, evolution should occur, but the size-related traits are in fact at evolutionary equilibria. Why does this happen? In short, the assumptions of additivity and the phenotypic gambit may be violated, the demographic functions that determine phenotypic selection may be incorrectly characterised, or the non-zero heritabilities may be an illusion. I argue here that, for size-related traits exhibiting stasis, the assumption of the phenotypic gambit is violated. However, to understand why, it is useful to address a question that is yet to be satisfactorily answered: why are organisms the size that they are (Smith and Lyons 2013, Bhat et al. 2020)?

Body size is a fundamental component of life history strategies. Within a species, individuals that are able to grow to large sizes are typically able to invest proportionally more energy in reproduction than those that grow to smaller sizes. Such within-species hyper-allometric scaling of reproductive investment is widespread, and may be universal (Barneche et al. 2018). Achieving large size is consequently desirable because it results in a fitness return via an increase in reproductive output. But if this is the case, why are not all species large? There is a risk to individuals of growing to large sizes before starting reproduction: development takes time, and delaying reproduction to older ages increases the risk of death before sexual maturity is reached (Blanckenhorn 2000). There is consequently a time to maturity-reproductive output trade-off that is determined by the associations of body size with survival, reproductive output, and development rate throughout the life course (Childs et al. 2004, Bhat et al. 2020). At some point on this trade-off, fitness will be maximised, and that determines size and age at sexual maturity. The optimal age and size at sexual maturity, and adult body size, is consequently determined by the shape of age- and size-specific fitness and transition functions (Kentie et al. 2020). Given this, the evolution of body size can be predicted by integrating across the entire life cycle assuming trade-offs are appropriately identified (Childs et al. 2004). Reproductive output can be divided into two other aspects of the life history strategy: offspring number and offspring size (Fleming and Gross 1990). These two quantities also trade-off against one another, with there being optimal values for both litter size and mean offspring size.

Rates of development are largely determined by the ability of individuals to acquire resources from the environment (Reznick et al. 2000). Individuals utilise the energy they obtain by allocating it to maintenance (staying alive and acquiring more resources), development, and reproduction (De Roos et al. 2003, Kooijman and Kooijman 2010, Smallegange et al. 2017). The way that resources are partitioned varies with age and size. The optimum body size and life history strategy are achieved by adopting a specific energy budget that partitions energy into each component of the life history at each age and size (Bhat et al. 2020). There is consequently a very close association between energy budgets and life history; this has led some authors to redefine fitness in terms of energy (Brown et al. 1993). Although at one level such an approach is appealing, it is inconsistent with standard evolutionary theory where selection, and fitness, are strictly defined in terms of numbers of descendants produced (Fisher 1930). Energy accrual needs to be linked to survival, development, reproduction, and inheritance to understand body size evolution (Bhat et al. 2020).

Within in a population there is usually one dominant cause of death that kills the largest proportion of individuals. However, individuals of different ages, or sizes, may experience different causes of death. Nonetheless, at the level of the population there is usually a primary cause of death associated with the limiting factor. Individuals within a population compete for energy in the presence of a limiting factor that determined the primary cause of death and failure to breed (Berryman 2004). For example, if prey (food) availability limits a predator population, individual predators compete over prey. In order to be competitive at predation, individuals need to express phenotypic traits that aid successful prey capture and consumption. Such traits could include camouflaged pelage, fast running speeds, weaponry associated with killing prey, and an ability to consume prey quickly. Selection is consequently expected to operate most strongly on phenotypic traits associated with prey capture and consumption as prey availability limits the population. In contrast, prey living in a predator-limited population are selected to acquire food resources without being predated. They experience selection for the accrual of enemy-free energy. In general, selection is expected to operate most strongly on traits associated with the accrual of energy given a population’s limiting factors and the primary causes of death and failures to breed.

Those individuals with phenotypic trait values that are well-matched to the environment in which they find themselves are better able to accrue resources than those with less optimal phenotypic trait values (Charlesworth 1989). They consequently grow faster, to larger age-specific sizes, have lower age-specific mortality rates, and higher rates of age-specific reproductive output. Those individuals that accrue most resources consequently perform well at all aspects of life compared to those that accrue fewer resources, and this heterogeneity in resource accrual between individuals can mask life-history trade-offs (Van Noordwijk and de Jong 1986, Metcalf 2016). A focus on resource accrual traits is consequently likely key to understanding how individuals obtain utilise energy in as an optimal way as possible to optimise life history and energy budgets, and maximise fitness (Bhat et al. 2020).

We can break down the accrual of resources into two separate processes: detection and acquisition. The senses, and the ability to process the information they reveal, are associated with the detection of resources, while many behaviors and some aspects of morphology are associated with their acquisition. The optimal set of phenotypic traits, and their values, associated with the detection and acquisition of resources will vary as a function of the population’s limiting factor and the primary causes of death and failures to breed.

Once resources are accrued, they need to be utilised. To do this, at each age and size, individuals need to allocate the resources they have accrued to maintenance, development, and reproduction (Kooijman and Kooijman 2010), in a manner that is expected to maximise their reproductive value via optimising the life history (Goodman 1982, Bhat et al. 2020). This is a hard-to-solve state-dependent decision-making problem that is constrained (McNamara and Houston 1996). Life history constraints mean that the total amount of energy an individual can allocate to a particular activity remains within bounds. For example, a long-lived iteroparous species with low reproductive effort, cannot suddenly switch strategies and invest all its energy into a single reproduction event, sacrificing its own survival. Energy budgets are plastic, but within limits that are set by past fluctuations in energy availability that populations have adapted to (DeWitt et al. 1998). Phenotypic traits associated with resource detection, acquisition and utilisation are likely subject to stabilising selection. This is because a trait value below a particular threshold will likely carry survival or reproductive costs as it makes it challenging to accrue resources, while an over-engineered trait carries a cost. For example, consider eye sight in Mexican cavefish. Vision can be critical for detecting predators, mates, and energy. Invest too little in developing vision, and the risk is an early death or a failure to find a mate. However, vision is expensive, commanding up to 15% of an individual’s energy budget (Moran et al. 2015). Visual acuity beyond being able to observe a mate, a predator, or food, is over-engineered and carries an energetic cost, and this is why the loss of cave fish vision in predator-free environments is adaptive. There is consequently a sweet spot, where vision is good enough to thrive, but not too good to be too expensive. Individuals with trait values at the sweet spot will acquire most resources and get the most out of them given the limiting environment, are expected to develop at a faster rate, and will have highest survival and reproductive rates, while those with smaller or larger values will achieve smaller sizes and have lower expected fitness (Figure 4).

**Figure 4.**
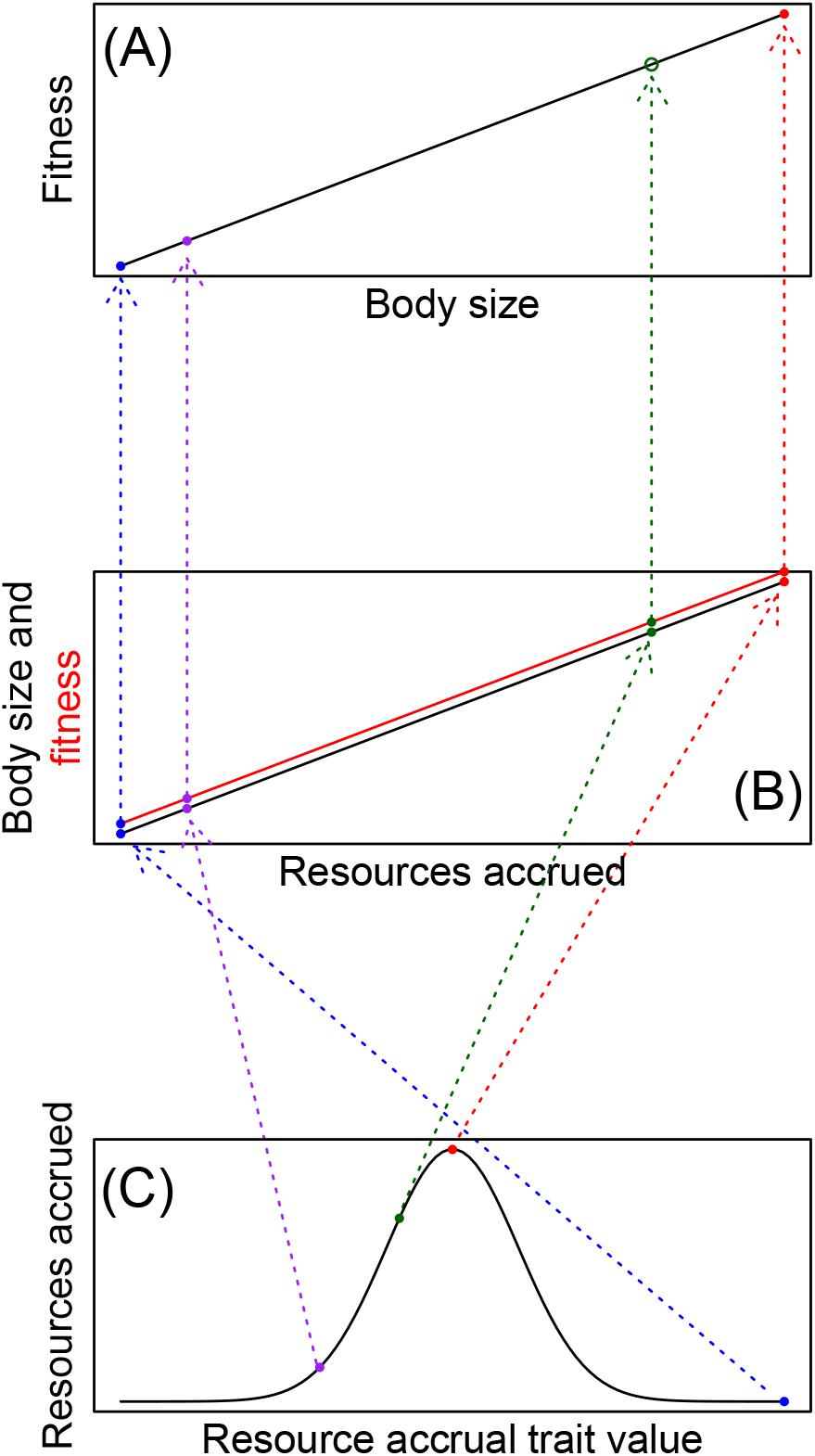
(A) A positive association between body size and fitness arises as (B) individuals that accrue most resources are larger and have higher fitness, which (C) is due to an n-shaped association between resource accrual and the resource accrual trait values. The dotted lines help visualize the map across functions.

If the resource accrual traits are heritable (and random mating is assumed), individuals with non-optimal resource accrual trait values and small body sizes will be expected to produce offspring that also have non-optimal trait values and small body sizes. In contrast, those individuals with resource accrual traits at the optimal value will be more likely to produce offspring that resemble them in these traits. Body size will consequently appear heritable and subject to directional selection (Figure 4), yet will not evolve to larger values because the mean of the resource accrual trait will be at the optimal value once the population has achieved evolutionary equilibrium. In addition, life history trade-offs will be hard to detect, because individuals with more optimal resource accrual traits and energy budgets acquire more energy and utilise it more effectively in development and reproduction than those with less optimal trait values (Van Noordwijk and de Jong 1986).

For the arguments above to hold, genetic variation needs to be maintained in the resource accrual trait. These traits may be determined by many loci of small effect, which means additive genetic variation is lost slowly (Lande 1982). Selection-mutation balance could theoretically contribute to the maintenance of genetic variation for resource accrual traits in such a scenario.

Frequency-dependent selection, operating via individual heterogeneity in resource use (Svanbäck and Bolnick 2006), could also maintain genetic and phenotypic variation via fitness-density covariance, stabilising mechanisms, the storage effect, or relative non-linearity (Bolnick et al. 2011). For example, if a new mutation generates a phenotypic innovation that allows an individual to utilise an under-utilised, or new, resource, that individual will accrue more resources, grow quickly, have high survival, and produce many offspring. If some of this individual’s descendants are also equipped to utilise the resource, the frequency of the new innovation will increase, but the novel resource being utilized will decline in abundance. Eventually the frequency of the new innovation will achieve an equilibrium point where it, and the resource, are in equilibrium. Random fluctuations in the biotic and abiotic environment can generate fluctuations in the numbers of genotypes and phenotypic traits, and this could generate frequency-dependent selection in resource accrual traits, contributing to maintain additive genetic variation.

It is possible that frequency-dependent and stabilising selection can operate simultaneously on resource accrual traits. If stabilising mechanisms contribute to the maintenance of genetic and phenotypic diversity, different phenotypic optima may exist among individuals that specialise on different resources. This would manifest as a flat-humped fitness function for the resource accrual trait when averaged across multiple time steps.

In this section I have argued that body size is determined by an interaction between breeding values *A_i_* and the environmental component of the phenotype *E_i_* such that body size is not determined by *Z_i_* ≠ *A_i_* + *E_i_* but rather by *Z_i_* = *A_i_* + *E_i_|A_i_* (Coulson et al. 2017). I propose that this is plausible, because *A_i_* is in fact a breeding value for a resource accrual trait subject to stabilising selection that impacts body size by determining the quantities of resources accrued by each individual. This conclusion is general: when a phenotypic trait appears heritable, is subject to directional selection, but does not evolve, then the assumption of additivity must be violated. Size-related traits appear to frequently violate this assumption. But in some cases, the amount of resources accrued may be determined by body size. How might this affect evolution?

## Body size as a resource accrual trait

In some instances, individuals with large body size are able to access more resources than those that are smaller. For example, large lions can prevent smaller members of the pack feeding on a kill, while large individuals of many species may be able to dominate smaller individuals in competition for a territory. Alternatively, it may be aggression that is the resource accrual trait determining the quantity of resources accrued, such that body size is a consequence of how aggressive an individual is, rather than body size per se being the resource accrual trait.

Body size also has the potential to play a key role as a resource accrual trait in seasonal environments where resource availability varies within a year as can be seen by returning to the long-run stochastic growth rate. The long run stochastic growth rate can be approximated as

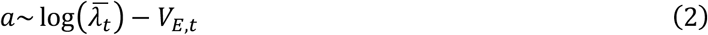

where the first term is the log of the mean of the per time-step population growth rates, and the second term, *V_E,t_*, is a function of the logged variance in the per time-step population growth rates (Lande et al. 2003). When *a* = 0, then obviously log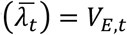. Mean fitness can consequently be increased when evolution operates on traits that either increase 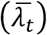 or decrease *V_E,t_* (Lande et al. 2017). Body size, hibernation, and migration are all traits that can evolve to decrease *V_E,t_* in seasonal environments. For example, individuals of large size of warm-blooded species have a higher body mass to volume ratio than their smaller counterparts and consequently use relatively less energy to keep warm in cold environments (Coulson et al. 2001). A consequence of this is that they may be able to survive winters more easily at higher latitudes than smaller individuals. Bigger individuals can also store larger fat reserves than smaller individuals to utilise during winter when energy is scarce. Large body size may consequently allow non-hibernating species to exist in areas where smaller competitor species cannot survive, thus accessing resources that are only available to them. However, the flip side to this, is that large size also requires more energy for maintenance (Brown et al. 1993, West et al. 1997), so resources must be sufficiently abundant during the summer months for individuals to accrue sufficient resources to lay down enough fat to survive the winter. It is possible that productivity in the growing season, and the harshness of the winter season, may both select for larger size within species as distance to the poles decreases, a phenomenon termed Bergmann’s rule (Bergmann 1848). It is also possible that climate change resulting in shorter and warmer winters size may allow smaller individuals to survive, leading to a reduction in body size within populations (Ozgul et al. 2009).

In some cases when body size is a resource accrual trait, it can conform to the additivity assumptions of quantitative genetics, and can evolve as predicted by the Breeder’s equation. Evidence for this comes from the observation that when body size is selected for in artificial settings it can evolve as predicted using methods from quantitative genetics. This insight might help explain why quantitative genetics so often works well in artificial environments but fails to do so in natural ones (Merilä et al. 2001b).

In artificial environments, energy is easy to detect and to acquire *ad libitum.* Selection is consequently largely relaxed on resource accrual traits, but is instead targeted at phenotypic traits we consider desirable. Only those individuals with desirable trait values are permitted to breed. This targeting selects for individuals that budget the largest proportions of the energy they accrue to the traits we desire. Artificial selection consequently selects for particular energy budgets that result in the traits we wish to evolve.

In the above sections I argue that populations and communities achieve ecological and evolutionary quasi-equilibria for protracted periods of time over which species composition remains constant. I then explain the set of conditions required for these equilibria to be attained. I now consider how models of the phenomena described above can be constructed.

## Population models of ecological and evolutionary stasis

In the section “**Characterising populations at ecological and evolutionary equilibrium**” I briefly explain that models can be structured to track the dynamics of distributions of individual attributes based on functions that describe associations of these attributes with survival, reproduction, development, and inheritance. I now expand on these earlier comments.

Structured demographic models can be constructed in discrete or continuous time. Discrete- and continuous-time modelling has proceeded largely independently, but has arrived at remarkably similar conclusions. Both approaches combine demographic, development, and inheritance functions to track the dynamics of distributions of genotypes and phenotypic traits (Figure 3). Models can be structured or individual-based (e.g. (Dunlap et al. 2006, Romero-Mujalli et al. 2019).

Discrete time models called integral projection models (IPMs) of phenotypic trait distributions were initially developed by (Easterling et al. 2000). Since then, models have been extended to incorporate explicit genotypephenotype maps (Coulson et al. 2011, Childs et al. 2016, Coulson et al. 2017, Rees and Ellner 2019), frequency-dependence (Schindler et al. 2015), density-dependence (Rees et al. 2014), abiotic variation (Ellner and Rees 2006), and species interactions (Adler et al. 2010, Bassar et al. 2017, Lachish et al. in press). These models have been used to study the dynamics of populations and distributions of phenotypic traits, breeding values and additive genotypes, non-additive genotypes, and life history strategies. The work has identified circumstances when eco-evolutionary feedbacks occur (Coulson et al. 2011). In these models, fitness, population dynamics and evolution are all emergent properties of the demographic, development and inheritance functions used to construct the models.

Continuous time models that track distributions of individual attributes have started to receive significant attention more recently. For example, (DeLong and Gibert 2016) introduced Gillespie eco-evolutionary models that are a form of continuous time individual-based model, and (Doebeli et al. 2017) argue that evolutionary models developed from birth and death rate functions to track distributions of heritable phenotypic traits are useful because fitness is an emergent property that does not need to be defined. Along similar lines, (Lowe et al. 2017) argue that demography and population genetics need to be explicitly combined to understand eco-evolutionary dynamics.

Because the two separate literatures have arrived at the same conclusions, it suggests that models based on demographic, development and inheritance functions offer great promise in providing novel ecological, evolutionary, and eco-evolutionary insight. I now focus on discrete time models that track distributions of alleles, genotypes, breeding values, and phenotypes for three reasons. First, the theory is more developed than for continuous time models, second, the approach has gained more traction amongst empiricists, and third, I am more familiar with this approach than with continuous time equivalents. However, the conclusions will extend to continuous time models.

IPMs are of the general canonical form:

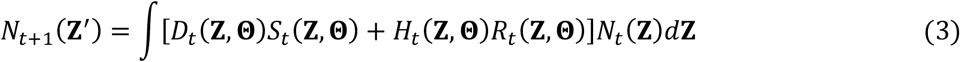

where **Z** is the set of individual attributes being iterated forwards, *N_t_*(**Z**) is the distribution of these attributes, the functions *S_t_*(**Z,Θ**) and *R_t_*(**Z, Θ**) are demographic functions describing how the individual attributes and the environment **Θ** at time *t* influence the rates of survival and reproduction respectively, and the functions *D_t_*(**Z.,Θ**) and *H_t_*(**Z,Θ**) are transition functions describing development and inheritance of the individual attributes. These functions are probability density functions such that they can move individuals between individual attribute classes but they do not alter the size (or volume) of the distribution. The transition functions can become quite complicated when different attributes included in Z have different development and inheritance rules (Schindler et al. 2013, Childs et al. 2016, Coulson et al. 2017). Feedbacks occur when properties of *N_t_*(**Z**) contribute to the environment **Θ** either directly or indirectly via interactions with other species.

How can models be constructed that achieve ecological and evolutionary equilibrium? First, these models need to iterate forward the dynamics of genotypes and phenotypic trait distributions. Demographic functions describe the growth rate of each part of the distribution being iterated but do not move individuals between categories; transition functions – e.g. development and inheritance functions – describe these movements (Coulson 2012). Second, models need to include ecological feedbacks such as species interactions, and stabilizing or frequency-dependent selection operating on genetic variation.

Demographic and transition functions are consequently central to constructing structured models of ecological and evolutionary dynamics and feedbacks. A key challenge in constructing models for multivariate distributions is that different types of individual attributes, or structural classes, have different development and inheritance properties, and consequently require different functional forms for their transition functions (Childs et al. 2016, Coulson et al. 2017) (Figure 3). Genotypes and breeding values, for example, remain fixed throughout life such that development functions simply keep individuals in the same class throughout life (Charlesworth 1994). For models containing these attributes to achieve a non-trivial evolutionary equilibrium, demographic functions need to capture a process such as frequency-dependent selection that maintains additive genetic variance, while simultaneously preventing directional evolution occurring indefinitely. Transition functions are also required to capture genetic inheritance, and the formation of offspring genotypes and breeding values as a function of parental values and patterns of mate choice (Figures 4(A-D)).

If I assume that resource accrual traits are additive and heritable and retain a fixed shape throughout life, but scale with body size, demographic and genetic inheritance functions that are similar to those for genotypes and breeding values are required, but so too is a development function that specifies how the size of the trait changes across the lifespan via changes to the environmental component of the phenotype (Figures 4(E-H)). These development functions keep the breeding values or genotypes associated with the resource accrual traits constant throughout life, while allowing the trait to develop with age, via change in the value of the environmental component of the phenotype.

Traits that exhibit the paradox of stasis such as size-related traits at equilibrium, need to develop with age, may show directional selection, but are not specified with an additive genotype-phenotype map (Coulson et al. 2010). Large parents may produce large offspring, and traits may statistically appear to be heritable but do not evolve (Figures 4(L)). Traits such as this require demographic, development, and inheritance functions, but the inheritance functions do not need to explicitly include rules for genetic inheritance for the trait itself because these traits are emergent consequences of an individual’s resource accrual trait value and the environment. Similarly, development functions can be specified to be determined by the value of resource accrual traits and the quantity of resources accrued (Lachish et al. in press).

The final trait type is life history traits such as size and age at sexual maturity. These traits are often referred to as latent traits, and they are only expressed once during an individual’s life (Childs et al. 2016). If these traits are under genetic control, the underlying genotypes need to be tracked throughout life, with the trait only expressed at one point in time, determining when reproduction occurs for the first time. The genetic architecture underpinning the trait can be subject to selection when the trait is not expressed if the trait genetically covaries with phenotypic traits that are expressed and are subject to selection.

IPMs have been developed to track the dynamics of each trait type, but all trait types are yet to be combined into a single model. One way to do this would be to construct a model of i) a resource accrual trait that is determined by a breeding value and an environmental component of the phenotype, ii) body size, where the development function is determined by the value of the resource accrual trait and the amount of resources accrued, and iii) age and size at sexual maturity is determined by a breeding value and an environmental component of the phenotype. An initial model might focus on a resource species and a consumer species. The resource accrual trait in the resource species would define the ability to escape consumption, while the resource accrual trait in the consumer species would define success at consuming resources.

Lachish *et al.* (in press) have developed a model of a vegetation resource that renews seasonally, and stochastically. The resource is consumed by elk (*Cervus canadensis*), with individuals in each attribute class partitioning the energy they accrue into maintenance, development, and, if individuals have reached sexual maturity, reproduction. A proportion of the maternal energy reserve of breeding individuals is invested into offspring. In this bioenergetic model based on dynamic energy budget theory (Kooijman and Kooijman 2010), the life history is determined by the energy budget at each age and size. This work builds on a previous model of Smallegange *et al.* (2017) that was the first to incorporate energy budgets into IPMs.

In such a model where the initial conditions are away from the evolutionary and ecological equilibrium, the breeding values for the resource accrual traits will change at each time step in a way determined by the covariance between breeding value and annual survival and reproductive rates at each age (Coulson et al. 2017). The dynamics of these traits will be accurately predicted by their per time-step selection differentials. Body sizes will change with time as the distribution of the resource accrual traits and the numbers of resources and consumers change. Body size dynamics will not necessarily be well-predicted by their per-time selection differentials. Neither will be the dynamics of age- and size at sexual maturity (Childs et al. 2011). In order to understand the dynamics of the resource accrual trait it is necessary to understand how evolution optimizes the trade-off between the risk of mortality prior to sexual maturity against any benefits of delaying reproduction to a later stage of development (Childs et al. 2004). The optimization of the trade-off maximizes reproductive value. A feedback now occurs: as reproductive value is maximized, age-specific survival and reproductive rates change, which means that the distribution of annual fitness changes, which alters the strength of selection on the resource accrual traits. However, eventually an ecological and evolutionary equilibrium will be achieved.

## Transitions between equilibria

Sudden shifts in the environment **Θ** be they biotic, such as the addition or removal of a species, or abiotic, such as a volcanic eruption, can alter the causes of death or failures to breed in some species. These changes will alter predicted values of the demographic and transition functions. For example, the loss of a predator species may result in a shift from top-down to bottom-up regulation of a prey species (Figure 5(A)-(B)) reflected by a reduction in the influence of a predator population on survival, reproduction, development or inheritance of a focal species, coupled with an increase in the role of a resource species on these functions. Such a change in a limiting factor can result in rapid change in population size of affected species (Figure 1(A)), and alter selection pressures on phenotypic traits associated with resource accrual (Figure 5(a)-(B)) and utilization (Figure 5(C)). Novel selection on these traits can result in a shift in phenotypic trait means via plasticity, (Figure 1(B)), and a slower rate of evolution, altering the frequency of alleles associated with resource accrual traits. Evolution occurs at a slower rate to the initial change in population size, such that the long-run stochastic growth rate of the population rapidly returns to being close to zero following the initial environmental perturbation. The novel selective regime on resource accrual traits can alter the optimal form of associations between body size and survival, reproduction, development, and non-genetic inheritance, leading to evolution of the life history. Change in the life history may result in additional novel selection pressures on resource accrual traits – a form of eco-evolutionary feedback.

**Figure 5:**
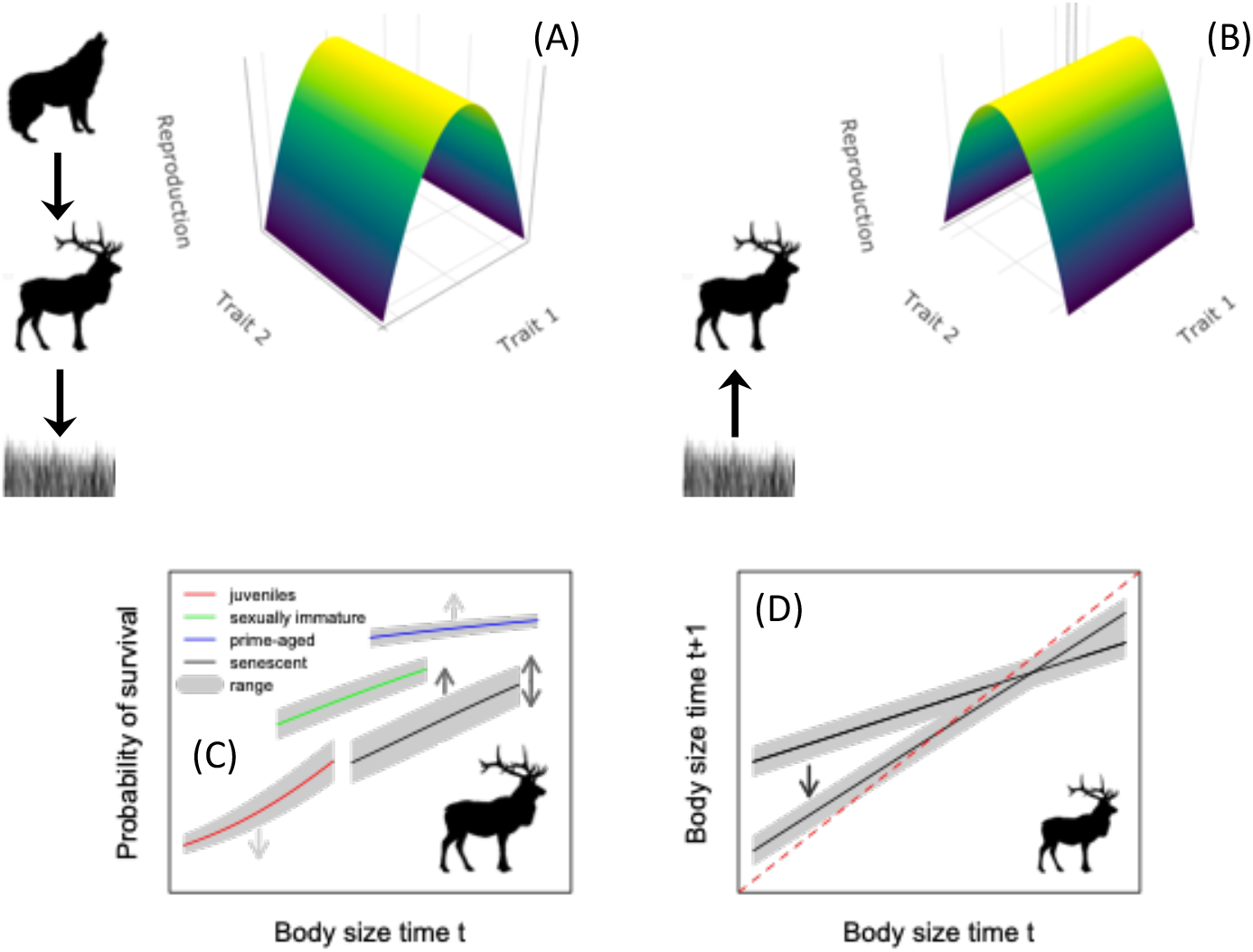
An environmental perturbation, such as the loss of a predator, can result in a period of transience between ecological and evolutionary equilibria. The removal of a predator can result in a change in limiting factor and a switch from top down (A) to bottom up (B) regulation. The change in the primary cause of death and failure to reproduce can relax selection on phenotypic traits associated with the original limiting factor and impose selection on traits associated with the new one (3D surfaces in A and B). These changes can alter demographic (C – survival as an example) and transition (D – development as an example) functions for size-related traits. In this example, a switch in limiting factor an increase in prime adult survival occurs with a simultaneous decrease in juvenile survival (C, gray arrows), an increase in the mean and variance in senescent survival (C black arrows), and a slowing in development rates (D, black arrow). In the original and novel equilibrium these functions take values that result a long stochastic growth rate of zero, although the mean and variance in population size may have changed (Figure 1(A)).

If a change in a limiting factor, population size, and selection pressures on resource accrual traits impacts the size of “keystone species” within the system, trophic cascades can occur where populations of other species within the ecosystem are impacted (Paine 1966, Bronstein et al. 2004, Georgelin et al. 2015). In this case, the distances between the population structures of the species within the community at the initial and new equilibrium will be large (Johansson and Dieckmann 2009). In contrast, if the changes in one species do not have substantial effects on keystone species, then a major shift in ecological and evolutionary equilibrium may not be observed (Brown and Vincent 1992).

The most pronounced trophic cascades occur when the composition of the dominant producer species changes. For example, in Yellowstone, it is argued that the decline in elk numbers due to increased wolf predation has led to forests replacing grasslands (Fortin et al. 2005). The effect is currently localized to small areas, as the two decades since wolves were reintroduced is not sufficient time for a forest to displace a grassland. However, if the effects observed on small spatial scales are eventually repeated on a larger scale, the reintroduction of wolves will be responsible for a pronounced trophic cascade as forests replace grasslands on the Northern range.

Less pronounced trophic cascades can occur when only a small part of a food web is affected, that does not impact primary producers. For example, a decline in the density of a predatory bird may influence the dynamics of its passerine prey and the insects on which they feed, but effects on the species structure of the primary producers is likely to be minimal (Borer et al. 2005). Those species directly impacted may experience altered causes of death or failures to reproduce, and novel selection pressures, but community structure and ecosystem function is unlikely to be significantly impacted to a sufficient extent to generate a pronounced trophic cascade.

Transitions between equilibria, whether they produced pronounced trophic cascades or not, can be captured with structured demographic models of interacting species that have achieved ecological and evolutionary stasis by setting the population of one species in the community to zero, by introducing a new species, or by changing the distribution of an abiotic driver. Any function that includes population size of the removed or added species, or the abiotic driver, will be directly impacted. Indirect effects will then cascade to other species whose demography might not be directly impacted by environmental perturbation, but might impacted by species that are directly impacted.

## Conclusions

The motivation for this paper was understanding the observations that i) communities often appear to be in equilibrium for protracted periods of time, yet ii) maintain additive genetic variance for many traits, allowing them iii) to evolve following an environmental perturbation, yet iv) many species exhibit traits, such as size, that do not evolve as predicted but instead exhibit stasis. When systems are perturbed from their equilibrium by the addition, or removal, of a species, they tend to a new ecological and evolutionary equilibrium (Bronstein et al. 2004, Georgelin et al. 2015). Individuals that are well-matched to their environment can exhibit different life history strategy strategies and resource accrual traits, or trait values, in each of the equilibrium states, with selection on these traits determined by the limiting environment. I have focused on traits associated with the accrual of food, but similar arguments hold for traits associated with the acquisition of mates. In order to understand observed dynamics, I have focused on the properties of ecological and evolutionary equilibria, the conditions required for these properties to be observed, how systems are expected to move from one state to another, and how one could go about modelling these dynamics using existing, well-developed, approaches (see also Johansson & Dieckmann 2009).

In this paper I have linked population limitation to community dynamics to evolution via biotic interactions and the accrual of resources. The ideas I present are consistent with many observations from a range of systems, but some are speculative. This manuscript should be seen as a perspective, rather than a comprehensive review. I hope that rather than being the last word, the paper generates discussion, tests of some of the hypotheses proposed, and acts to stimulate new models of the interplay between ecology and evolution.

## Acknowledgements

Version 4 of this preprint has been peer-reviewed and recommended by Peer Community In Ecology (https://doi.org/10.24072/pci.ecology.100053)

The ideas in this paper have arisen from numerous discussions and arguments with friends, colleagues, and adversaries over the years. In the friend and colleague camp I would like to thank Ron Bassar, Tim Benton, Ottar Bjornstad, Mark Boyce, Sonya Clegg, Tim Clutton-Brock, Sasha Dall, Andy Dobson, Jean-Michel Gaillard, Alan Grafen, Bryan Grenfell, Pete Hudson, Bruce Kendall, Loeske Kruuk, Virpi Lummaa, Dan MacNulty, Dan Nussey, Josephine Pemberton, David Reznick, Andy Russell, Bernt-Erik Saether, Isabel Smallegange, Doug Smith, Dan Stahler, Joe Travis, Shripad Tuljapurkar, Alastair Wilson and all past and present members of my research group, for stimulating science discussions. I also owe a debt of gratitude to Marco Festa-Bianchet, Michael Morrissey, Erik Postma, and Norman Owen-Smith who I have not always agreed with, but whose critiques influenced my thinking; thank you, and apologies if our arguments caused any stress. I would also like to thank NERC, BBSRC, the Wellcome Trust, and the ERC for funding. Having been funded to work on systems as diverse as the guppies of Trinidad (thanks Ron, Joe and David), the wolves of Yellowstone (thanks Doug, Dan and Dan), the Soay sheep and red deer of Scotland (thanks Tim CB, Josephine, Dan N, Alastair, and Loeske), and the mites of Holland (thanks Isabel) is a real privilege, and each system has contributed to the ideas described here.

